# Modulating the transcriptional landscape of SARS-CoV-2 as an effective method for developing antiviral compounds

**DOI:** 10.1101/2020.07.12.199687

**Authors:** Daisy A. Hoagland, Daniel J.B. Clarke, Rasmus Møller, Yuling Han, Liuliu Yang, Megan L. Wojciechowicz, Alexander Lachmann, Kasopefoluwa Y. Oguntuyo, Christian Stevens, Benhur Lee, Shuibing Chen, Avi Ma’ayan, Benjamin R tenOever

## Abstract

To interfere with the biology of SARS-CoV-2, the virus responsible for the COVID-19 pandemic, we focused on restoring the transcriptional response induced by infection. Utilizing expression patterns of SARS-CoV-2-infected cells, we identified a region in gene expression space that was unique to virus infection and inversely proportional to the transcriptional footprint of known compounds characterized in the Library of Integrated Network-based Cellular Signatures. Here we demonstrate the successful identification of compounds that display efficacy in blocking SARS-CoV-2 replication based on their ability to counteract the virus-induced transcriptional landscape. These compounds were found to potently reduce viral load despite having no impact on viral entry or modulation of the host antiviral response in the absence of virus. RNA-Seq profiling implicated the induction of the cholesterol biosynthesis pathway as the underlying mechanism of inhibition and suggested that targeting this aspect of host biology may significantly reduce SARS-CoV-2 viral load.

## Introduction

A zoonotic transmission of an airborne virus capable of rapid spread has always been a potentially devastating threat to human society; this risk has only increased with globalization. Indeed, the recent emergence of SARS-CoV-2 has proved an unprecedented global challenge. As the scientific community works fastidiously to learn more about the biology of the virus and develop a vaccine, there is a pressing need to identify therapeutics to reduce morbidity and mortality. To better understand the basis of the disease, initial efforts have focused on determining how different cell types and tissues respond to SARS-CoV-2 infection (Blanco-Melo et al., 2020; Zhou et al., 2020; Ziegler et al., 2020). These efforts have generated a high-level understanding of how virus replication correlates with induction of the antiviral response and the changes in cellular processes. In general, a viral infection results in the production of numerous inflammatory triggers which can include aberrant RNA, misfolded proteins, and/or damage to the cell (Devarkar et al., 2016). In the case of SARS-CoV-2, the virus appears to mask aberrant RNA to prevent a robust induction of the cellular antiviral response. However, the processes of SARS-CoV-2 replication still result in a unique transcriptional footprint that presumably accommodates the virus life cycle (Blanco-Melo et al., 2020). Here we attempt to leverage this information to predict compounds that counter these conditions by inverting transcriptional signatures.

Ideally, in the face of a pandemic we would be able to deploy a pan-antiviral that universally targets a transcriptional footprint common to all viruses. As viruses are obligate parasites, they almost exclusively require host machinery to effectively replicate and spread, making it difficult to develop an antiviral nontoxic to the host. Adding to this challenge is the fact that every human pathogen antagonizes some portion of the cellular antiviral response making each transcriptional landscape unique (García-Sastre, 2017). While development of family-specific antivirals has had success with viruses such as orthomyxoviridae (Krammer et al., 2018). and human immunodeficiency virus (Gulick and Flexner, 2019), such antivirals still do not exist for the coronavirus family. In the absence of a pan-specific coronavirus drug, or a SARS-CoV-2 vaccine, the next most useful tool would be the effective repurposing of FDA-approved drugs. As this expansive list of compounds has been characterized in humans for toxicity and delivery, identified compounds could be immediately deployed should they show beneficial effects. So far, most drug repurposing efforts involve computational predictions via structural biology, network analysis, or *in vitro* drug screens (Gordon et al., 2020). Until now, these drug screens have produced low overlap (Kuleshov et al., 2020), a phenomenon likely due to many variables including the screen’s readout or the choice of cell platform. There is underutilized opportunity to understand the global molecular changes these drugs induce by using available sequencing data from diverse models and cell types.

The library of integrated network-based cellular signatures (LINCS) is an NIH consortium working to define multi-omics footprints for pre-clinical and approved drugs (Keenan et al., 2018; Subramanian et al., 2017). This resource serves as a promising computational method to find compounds which may counter or mimic the gene expression changes induced by a given perturbation. By comparing changes in gene expression before and after drug treatment, and/or viral infection, we can identify transcriptional irregularities that are inversely correlated between these two groups. Applied here, such an approach may methodically identify antivirals whose mechanisms of action are based in returning cells to homeostasis while impairing virus replication. As past efforts successfully utilized the LINCS resource to identify drugs that attenuate Ebola virus in cell-based assays (Duan et al., 2016), here we applied this approach to identify antivirals for SARS-CoV-2. By analyzing expression profiling of SARS-CoV-2 infected cells, we identified a high-dimensional region in expression space unique to this virus, namely cholesterol biosynthesis, whose perturbation interferes with virus biology. These drugs were validated in multiple cell-based assays including SARS-CoV-2-infected Vero cells, A549^ACE2^ cells, and in human organoids. In all, these efforts identified four promising drug candidates: amlodipine, loperamide, terfenadine and berbamine which each inhibit the virus independent of viral entry.

## Results

In a recent study (Blanco-Melo et al., 2020), we applied gene expression profiling to generate transcriptional signatures from lung cultures infected with SARS-CoV-2. Those analyses revealed that virus infection resulted in robust replication and a strong cellular response. As a follow up, we used the RNA-seq data collected from these studies and processed them using BioJupies (Torre et al., 2018) to create differential expression signatures (see methods). These signatures were then used as queries against the LINCS L1000 dataset, a collection of gene expression profiles generated following the administration of >20,000 bioactive compounds including >1,000 FDA-approved drugs to human cell lines at a variety of different times and concentrations (Subramanian et al., 2017) With L1000FWD (Wang et al., 2018), we could identify reciprocal transcriptional signatures generated between SARS-CoV-2 infection and a given compound. Specifically, we identified a common region in the L1000FWD expression space that is not well characterized, and where genes down-regulated by SARS-CoV-2 infected cells are up-regulated by a collection of drugs and small molecules profiled by the L1000 platform. By manually examining the drugs that are consistently targeting this region in expression space, we identified: terfenadine, loperamide, berbamine, trifluoperazine, amlodipine, RS-504393, and chlorpromazine as having potential therapeutic value against SARS-CoV-2 (Figure 1A). In support of these findings, it is noteworthy that loperamide, berbamine, and trifluoperazine were recently reported as hits in an independent drug screen for compounds that block SARS-CoV-2 infection in African Green Monkey kidney (Vero-E6) cells (Jeon et al., 2020).

**Figure 1.**
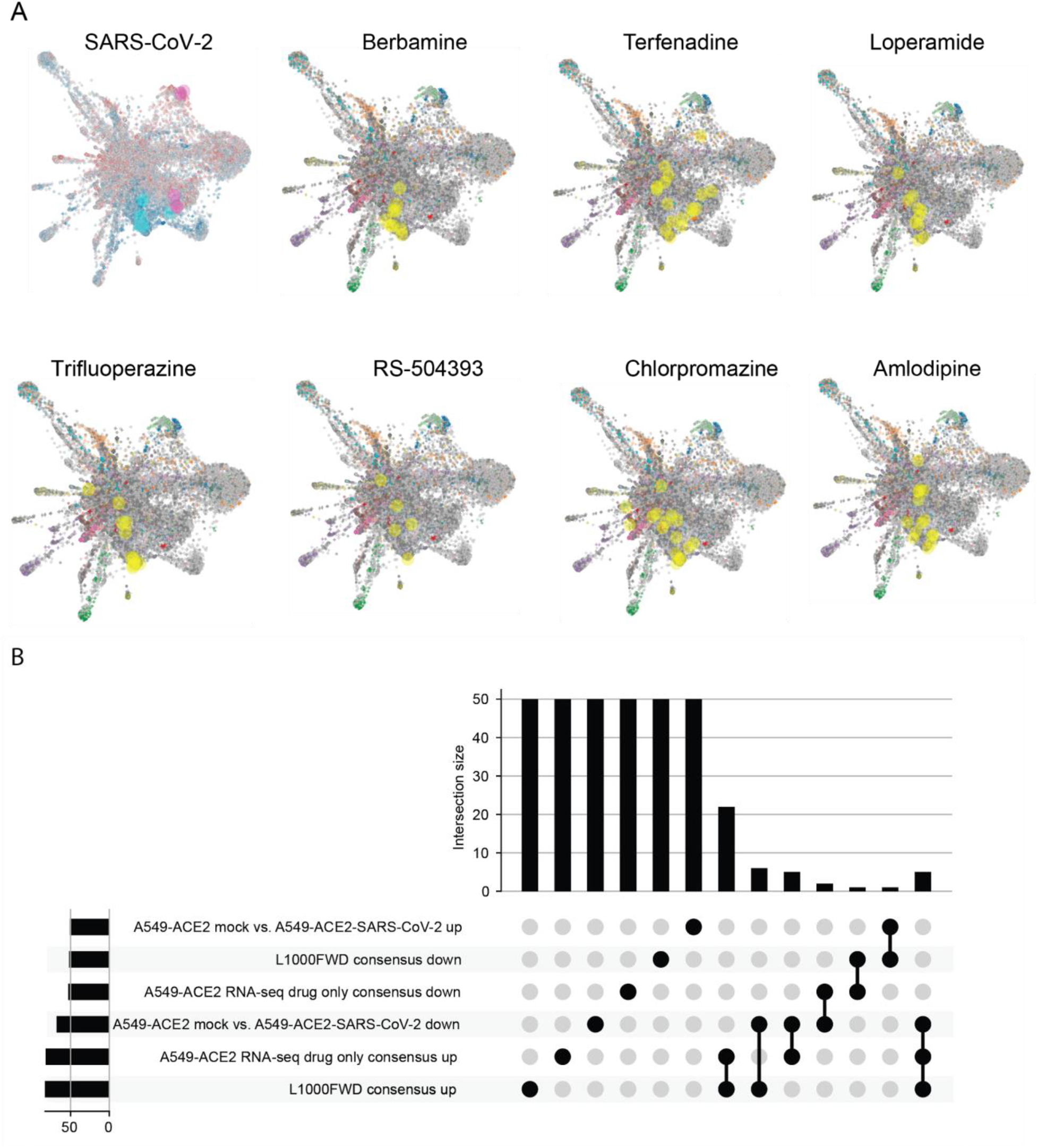
L1000FWD analysis of SARS-CoV-2 and candidate drug transcriptional space. A) L1000FWD analysis projecting the SARS-COV-2 uninfected vs. infected A549-ACE2 RNA-seq signature onto the L1000FWD space (a) where red represents mimicking signatures and blue represents reversing signatures. Top 5 mimickers are highlighted in pink and top 5 reversers are highlighted in cyan. Projection of 7 reversing drugs: terfenadine, loperamide, berbamine, amlodipine, trifluoperazine, RS-504393, and chlorpromazine. Here signatures are colored by their known mechanisms of action where the signatures for each drug are highlighted in yellow. B) UpSet plot to visualize the overlap between consensus top 50 up and down genes for the 6 drugs tested in A549-ACE2 cells by RNA-seq and from their collective signatures from the L1000FWD resource. These signatures are compared to the top 50 up and down genes computed from a signature created from SARS-CoV-2 uninfected vs. infected A549^ACE2^ cells from GSE74235. Significance overlap is observed for the intersection between the consensus up genes from the RNA-seq and L1000 drug only sets (22 genes, p-value<2.1e-46, Fisher exact test), and between the consensus up genes from the RNA-seq, L1000 drug only sets, and the top 50 down genes from the A549-ACE2 infected cells (5 genes, p-value<5.1e-101, empirical sampling fitted to a Poisson distribution). These 5 genes are listed above their associated bar.

The transcriptional signature implicating the aforementioned compounds include 50 genes down-regulated by the virus that are consistently up-regulated by these drugs (22 out of 50/50, p-value<2.1e-46, Fisher exact test; and 5 out of 50/50/50, p-value<1.7e-20, empirical simulations) (Figure 1B). Overall, based on the L1000 data, these seven compounds influence the same pharmacological high-dimensional gene expression signature space and are predicted to disrupt key cellular processes that are modulated in response to SARS-CoV-2 infection. Moreover, the expression signatures induced by all seven drugs across multiple cell-line contexts are highly correlated (Figure S1A). In addition to leveraging the L1000 data, we also examined the top 100 genes co-expressed with ACE2, the established receptor of SARS-CoV-2 (Hoffmann et al., 2020), based on correlations computed across thousands of diverse RNA-seq datasets (Lachmann et al., 2018). We then searched for FDA-approved compounds that induce or suppress the expression of these genes using drug signatures extracted from other independent transcriptome studies (Wang et al., 2016). This approach identified one additional compound, quercetin, as an effector of these ACE2 co-expressed genes (Figure S1B). Hence, all together, eight drugs were selected to be tested for efficacy against SARS-CoV-2 *in vitro*.

Next, we examined whether these eight drugs influenced SARS-CoV-2 infection of Vero-E6 cells by applying them at the same concentration used in the L1000 experiments (10μM) with the exception of quercetin (40μM), which was determined based on the concentration used in the top matching signatures extracted from GEO (Mutch et al., 2006). Out of the eight drugs, seven drugs completely abolished the presence of SARS-CoV-2 nucleocapsid when administered one hour prior to infection (Figure 2A). Quercetin was the only drug where the viral nucleocapsid remained present, while chlorpromazine induced cytotoxicity and therefore was not characterized further (Figure S2A). We next tested the remaining seven drugs in more relevant cell-based models. Specifically, we administered the compounds to an adenocarcinomic human alveolar basal epithelial (A549) cell line that constitutively expresses ACE2 (herein referred to as A549^ACE2^). We next pre-treated these cells with terfenadine, loperamide, berbamine, trifluoperazine, amlodipine, and RS-504393 and infected cells with SARS-CoV-2. In agreement with our Vero-E6 results, these data demonstrated that all six drugs successfully blocked virus replication (Figure S2B). These results could be further corroborated at an RNA level by performing quantitative RT-PCR based on the 5’ transcription-regulatory sequence (TRS) and the 3’ end of the N open reading frame which confirmed the effect of the six remaining drugs in both Vero-E6 (Figure S2C) and in A549^ACE2^ cells (Figure 2B). Lastly, to assess how these compounds impact viral growth, we performed plaque assay from the supernatant of treated A549^ACE2^ cells and found that terfenadine and berbamine completely inhibited the production of progeny virions below the limit of detection whereas loperamide, trifluoperazine, amlodipine, and RS-504393 all significantly reduced viral output by 2-3 orders of magnitude (Figure 2C). To determine if inhibition was mediated by a block in viral entry, we tested each compound for the capacity to block Renilla luciferase activity delivered by a recombinant Vesicular Stomatitis Virus pseudotyped with SARS-CoV-2 Spike. These data found that entry was not impacted in response to any of the tested compounds (Figure 2D).

**Figure 2.**
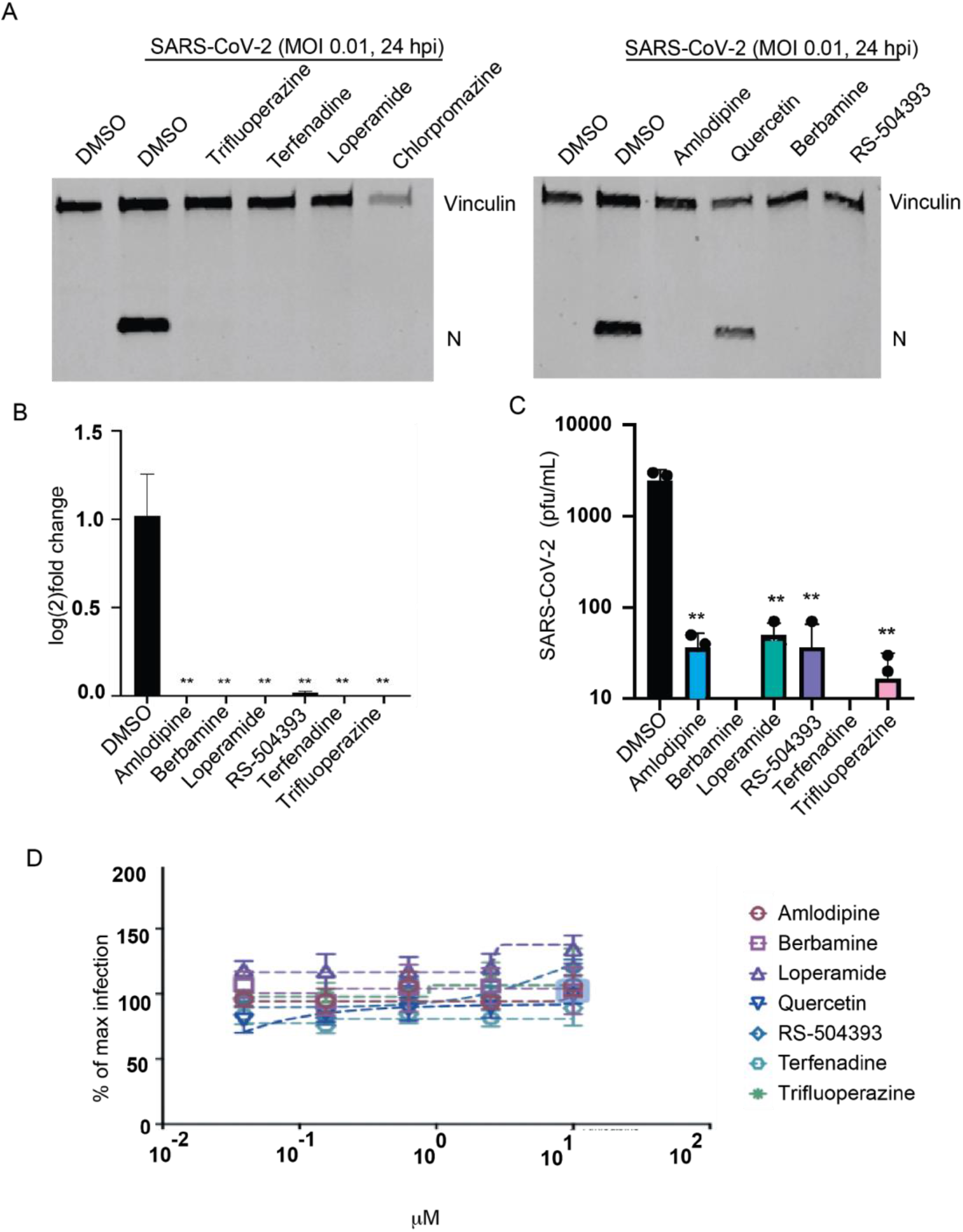
Drug screen in Vero-E6 and A549^ACE2^ cells. A) Western blots of SARS-CoV-2 infected Vero-E6 cells in presence of the eight predicted drugs; SARS-CoV-2 infection at MOI 0.01 for 24 hours. B) viral sgRNA quantification of SARS-CoV-2 infected A549^ACE2^ cells by qRT-PCR using TRS-L and TRS-N primers normalized to alpha tubulin levels; SARS-CoV-2 infection at MOI 0.1 for 24 hours. C) Viral titers of SARS-CoV-2 infected A549^ACE2^ supernatant treated with the six drugs that were not toxic and show promise in Vero-E6 cells. D) Entry inhibition assay of SARS-CoV-2 pseudotyped VSV particles measured by Renilla luciferase activity. Significance compared to DMSO treated values and determined by unpaired two-tailed student’s t-test, p<0.01 = **, n=3 biological replicates for qRT-qPCR and titer data.

To further understand the molecular basis for the inhibition of SARS-CoV-2, we next administered the drugs to A549^ACE2^ cells both in the absence and presence of virus and measured genome-wide gene expression with RNA-seq under all these conditions. As expected, in the absence of virus we observe that the six drugs induce expression signatures comparable to those generated by processing the L1000 data (Figure 3A, Figure S3A, and Tables S1-13). STRING annotation prediction software of differentially expressed genes (DEGs) demonstrated overlapping annotations between different drug treatments including the induction of cholesterol biosynthesis (GO:0006695), amino acid biosynthesis (GO:1901607), protein metabolism (GO: 1903364), and a downregulation of cell cycle (GO: 0007049) (Figure 3C). These transcriptional changes are also evident in the presence of virus infection and can be viewed individually on the corresponding volcano plots (Figure 3C-I). These data agree with qRT-PCR and Western blot analysis (Figure 2 and Figure S2). Moreover, SARS-CoV-2 viral reads per million reads (RPM) in the RNA-seq data are 3 magnitudes lower in the six drug-treated compared to DMSO-treated A549^ACE2^ populations (Figure 3B). Taken together, these annotation signatures implicate cholesterol biosynthesis and cell cycle as the two most prominent pathways modulated by the administration of these compounds. However, as there are thousands of other L1000 compounds known to impact cell cycle, these data would implicate cholesterol biosynthesis as the predominant mode of action in inhibiting SARS-CoV-2 in these assays.

**Figure 3.**
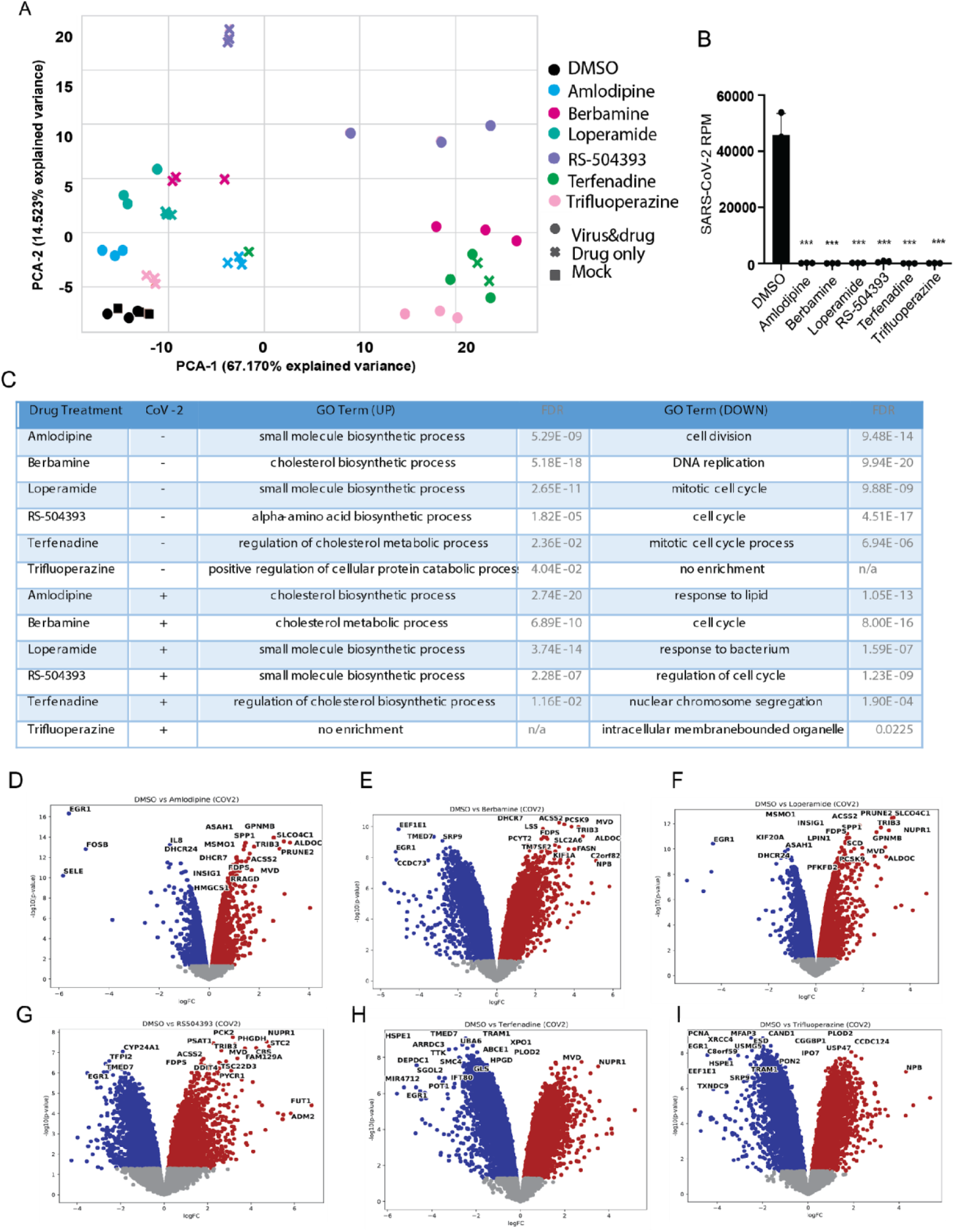
RNA-seq data analysis from A549^ACE2^ cells. A549^ACE2^ were either mock-infected (MOCK) and treated with DMSO or infected with SARS-CoV-2 in the presence of DMSO or the 6 indicated drugs for 24 hours at MOI 0.1. A) Principal component analysis of the RNA-seq data samples in PC1/PC2 space. B) SARS-CoV-2 reads in the RNA-seq data normalized to total reads per million of each sample. C) Top enriched GO term after STRINGdb analysis comparing either drug treated to DMSO-treated ACE2-A549s without infection, or drug-treated to DMSO-treated A549^ACE2^ after SARS-CoV-2 infection. D-I) Volcano plots of the infected A549^ACE2^ populations comparing DMSO- or drug candidate-treated after SARS-CoV-2 infection. Significance compared to DMSO treated values and determined by unpaired two-tailed student’s t-test, p<0.001=***; n=3 biological replicates for RNA-seq data.

To further assess the global effects of these drugs, we next applied them to human pluripotent stem cell-derived pancreatic endocrine organoid cultures, previously shown to support robust SARS-CoV-2 replication (Yang et al., 2020). In addition to testing the drugs for their ability to attenuate infection, we also performed genome-wide RNA-seq profiling to examine the effect of these drugs on the human pancreatic organoid cultures. Principal component analysis of infected organoids found that all these samples formed three major clusters in this transcriptional space defined by variance one and two (Figure 4A). These data demonstrate that PCA-1 could be largely defined as an uninfected vs. infected signature. One exception to this was berbamine, which formed a unique cluster for both variance one and two. When this same data was mapped on variance two and three, one could observe greater separation between the non-berbamine drug treatment and DMSO – reflecting the changes to cholesterol biosynthesis (Figure S4A).

**Figure 4.**
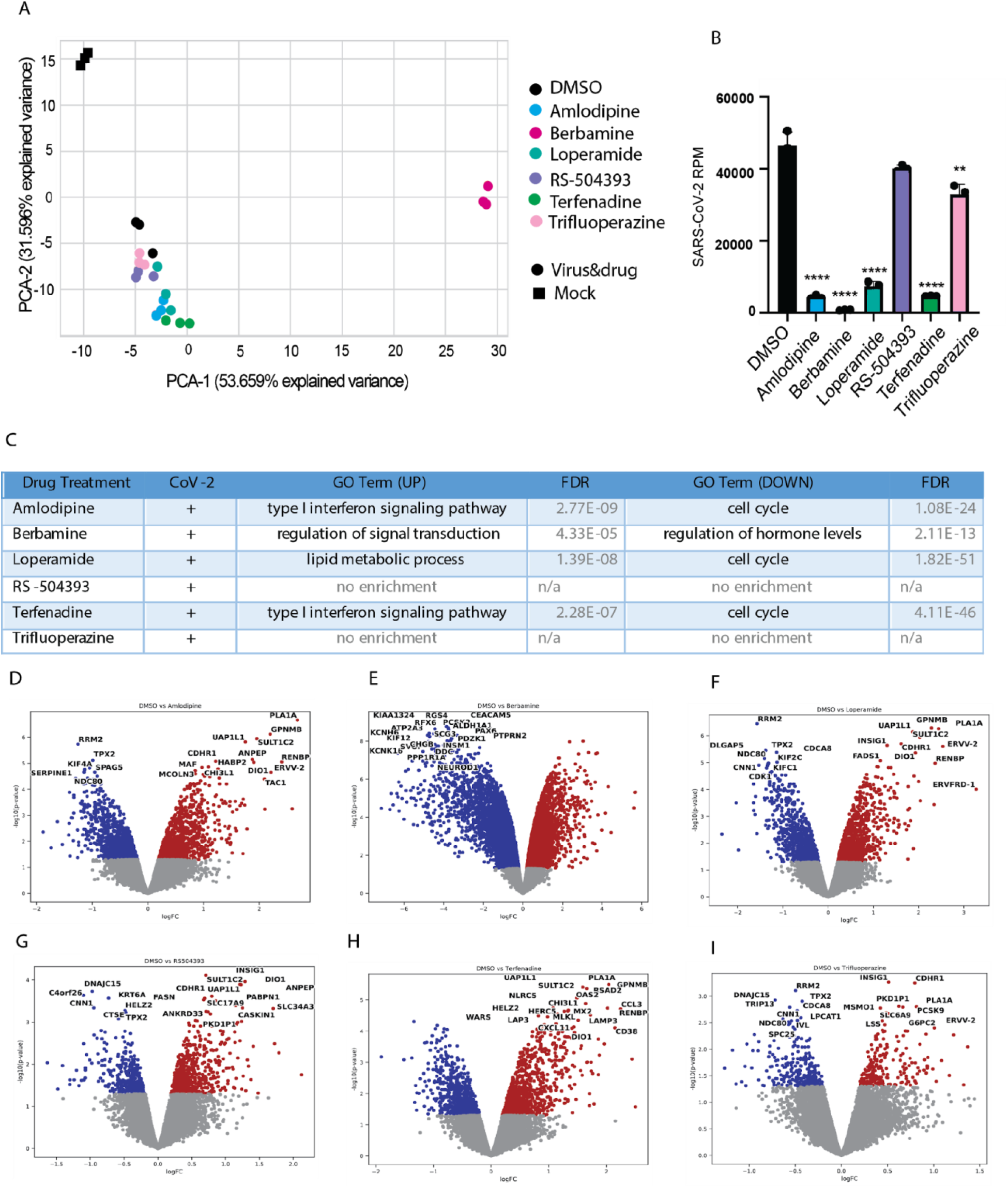
RNA-seq data analysis from pancreatic endocrine organoids. Human organoids were either mock-infected (MOCK) and treated with DMSO or infected with SARS-CoV-2 in the presence of DMSO or the 6 indicated drugs for 24 hours at MOI 0.1. A) Principal component analysis of the RNA-seq data samples in PC1/PC2 space. B) SARS-CoV-2 reads in the RNA-seq data normalized to total reads per million of each sample. C) Top enriched GO term after STRINGdb analysis comparing drug-treated to DMSO-treated human organoids after SARS-CoV-2 infection. D-I) Volcano plots of the infected human organoids comparing DMSO- or drug candidate-treated after SARS-CoV-2 infection. Significance compared to DMSO treated values and determined by unpaired student’s t-test, p<0.01 = **, p<0.0001 = ****, n=3 biological replicates for RNA-seq data.

Aligning captured reads to a SARS-CoV-2 reference genome, we find that while trifluoperazine and RS-504393 demonstrated only modest reduction of viral load, loperamide, terfenadine, amlodipine, and berbamine all successfully reduced virus transcription by approximately three orders of magnitude (Figure 4B). While genes involved in the cholesterol biosynthesis pathway are still upregulated in DEG analysis (Figure 4D-I, Tables S14-20), the most upregulated GO annotation in amlodipine and terfenadine is the type I interferon signaling pathway, and in berbamine treated human organoids is the regulation of signal transduction (Figure 4C), indicating a more nuanced mechanism of action in this more biologically relevant tissue model. Trifluoperazine and RS-504393 treated organoids’ lack of enrichment is reflected in decreased efficacy to lower viral reads (Figure 4B).

Taken together, these data suggest that the small molecules capable of reversing the transcriptional landscape induced by SARS-CoV-2 infection may be a *bona fide* means of identifying novel antiviral therapeutics. While additional *in vivo* testing will be required to ascertain whether amlodipine, berbamine, loperamide, and/or terfenadine represent effective antivirals against SARS-CoV-2, the findings outlined here also provide us with a greater understanding of virus-host interactions.

## Discussion

Here we identified several potential drug repurposing candidates based on analysis of published RNA-seq and L1000 gene expression data. Specifically, we identified a region in the pharmacological gene expression space that regulates cholesterol biosynthesis that appears to negatively impact virus replication. This led us to predict seven drugs as potential SARS-CoV-2 inhibitor candidates: terfenadine, loperamide, berbamine, trifluoperazine, amlodipine, RS-504393, and chlorpromazine. Loperamide is an over the counter approved drug used to treat diarrhea which is one of the symptoms of COVID-19. Berbamine is an experimental drug extracted from a shrub native to Japan, Korea, and parts of China (Greathouse and Rigler, 1940). It is an alkaloid and a channel blocker that was previously shown to display anti-lung-cancer (Duan et al., 2010) and anti-inflammatory activity (Jia et al., 2017). Trifluoperazine was shown to block Epstein-Barr virus (Nemerow and Cooper, 1984), while amlodipine is an approved drug for blood pressure and chest pain (angina) (Chahine et al., 1993). Amlodipine was recently reported to improve the outcome of patients with COVID-19 (Solaimanzadeh, 2020; Zhang et al., 2020). The preclinical drug RS-504393 is a known CCR2 antagonist that blocks CCL2 signaling in sepsis (Souto et al., 2011). Hence, its modulation of the immune response may interfere with SARS-CoV-2 host interactions. Terfenadine was withdrawn from the market in 1997 after 12 years due to evidence for cardiac toxicity via the binding to the hERG channel yet may still provide short term benefit as a SARS-CoV-2 prophylactic or acute treatment (Roy et al., 1996). Chlorpromazine was shown to block Epstein-Barr virus (Nemerow and Cooper, 1984) and influenza A virus (Krizanova et al., 1982) and very recently was found to be effective against SARS-CoV-2 in multiple in-vitro assays (Plaze et al., 2020), although we ruled it out early due to cellular toxicity.

These seven drugs were first tested in Vero cells by two independent cell-based assays with six of them showing positive results. The six most promising compounds were then tested in A549^ACE2^ cells and in pancreatic endocrine human organoids. The six drugs significantly diminished SARS-CoV-2 infection in A549^ACE2^ cells, and four out of six of these drugs also inhibited the virus in the human organoid model. RNA-seq profiling of A549^ACE2^ cells and the organoids in the presence of these drugs showed that berbamine displays quite different transcriptional effects compared with the other compounds. It has a more pronounced effect that is opposite in expression space, bringing the tissue closer to its normal uninfected state. It should be noted that the module of genes that are up regulated by the identified compounds is known to be targeted by commonly used cholesterol lowering drugs. It was suggested that statins may be considered as a potential treatment (Castiglione et al., 2020), but there is some evidence that statins may increase COVID-19 infection (Shrestha, 2020). While such effects need to be further determined, our analysis suggests that the same pathways are likely involved in SARS-CoV-2 pathogenesis and have potential for SARS-CoV-2 inhibition. In this regard it is noteworthy that SARS-CoV-2 replication demands extensive intracellular membranes (Snijder et al., 2020) which would be impacted by changes to cholesterol biogenesis, as is the case for Hepatitis C Virus, another positive sense RNA virus (Paul et al., 2013; Stoeck et al., 2018). While we demonstrated here the efficacy for several promising repurposed drugs as potential COVID-19 treatments in multiple cell-based assays, it remains to be determined whether these drugs will show any benefit to patients. The next step will be to test these compounds in an animal model before initiating clinical trials. Furthermore, with varying efficacy in different cell culture models and tentative value as therapeutics, all six aforementioned drugs clearly inhibit SARS-CoV-2 replication *in vitro*. Should animal models prove these drugs to be ineffective, these data would still be of value as they can be used to study the biology of SARS-CoV-2 as it pertains to cholesterol biosynthesis.

## Methods

### Cell Culture

Human adenocarcinomic alveolar basal epithelial (A549) cells (ATCC, CCL-185), African green monkey kidney epithelial Vero-E6 cells (ATCC, CRL-1586) were maintained at 37°C and 5% CO2 in Dulbecco’s Modified Eagle Medium (DMEM, Gibco) supplemented with 10% Fetal Bovine Serum (FBS, Corning). A549^ACE2^ heterogeneous cell population was generated by transducing A549 cells with lentivirus without selection. All drug treatments were administered to plated cells and cell cultures 1 hour prior to infection.

### Viruses

SARS-related coronavirus 2 (SARS-CoV-2), Isolate USA-WA1/2020 (NR-52281) was deposited by the Center for Disease Control and Prevention and obtained through BEI Resources, NIAID, NIH. SARS-CoV-2 was propagated in Vero-E6 cells in DMEM supplemented with 2% FBS, 4.5 g/L D-glucose, 4 mM L-glutamine, 10 mM Non-Essential Amino Acids, 1 mM Sodium Pyruvate and 10 mM HEPES. Infectious titers of SARS-CoV-2 were determined by plaque assay in Vero E6 cells in Minimum Essential Media supplemented with 4 mM L-glutamine, 0.2% BSA, 10 mM HEPES and 0.12% NaHCO3 and 0.7% agar. All work involving live SARS-CoV-2 was performed in the CDC/USDA-approved BSL-3 facility of the Global Health and Emerging Pathogens Institute at the Icahn School of Medicine at Mount Sinai in accordance with institutional biosafety requirements.

### Quantitative real-time PCR analysis

RNA was reverse transcribed into cDNA using SuperScript II Reverse Transcriptase (Thermo Fisher) with oligo d(T) primers. Quantitative real-time PCRs were performed using KAPA SYBR FAST qPCR Master Mix Kit (Kapa Biosystems) on a LightCycler 480 Instrument II (Roche). Primers specific for the leader sequence and nucleocapsid sgRNA (Primers: CTCTTGTAGATCTGTTCTCTAAACGAAC, GGTCCACCAAACGTAATGCG) and human alpha-Tubulin were used (Primers: GCCTGGACCACAAGTTTGAC, TGAAATTCTGGGAGCATGAC). Viral RNA levels were quantified by normalizing vRNA delta-Ct values to alpha-tubulin and calculating log(2)fold reduction normalized to DMSO-treated and infected samples. Significance was determined using an unpaired two-tailed student’s t-test.

### Western blot analysis

Protein was extracted from cells using Radioimmunoprecipitation assay (RIPA) lysis buffer containing 1X cOmplete Protease InhibitorCocktail (Roche) and 1X Phenylmethylsulfonyl fluoride (Sigma Aldrich) prior to removal from the BSL-3 facility. Samples were normalized for protein content using BioRad Protein Assay Dye Reagent (BioRad, 5000006) and run by SDS-PAGE electrophoresis. Proteins were transferred to nitrocellulose membranes and blocked in 5% milk in PBS. Proteins were detected using mouse monoclonal anti-SARS-CoV-2 Nucleocapsid [1C7C7] protein (a kind gift by Dr. T. Moran, Center for Therapeutic Antibody Discovery at the Icahn School of Medicine at Mount Sinai) and rabbit monoclonal anti-vinculin (Abcam, ab129002). Primary antibodies were detected by IRDye 680 Goat anti-Mouse IgG secondary antibody (926-68070) and IRDye 800 Goat anti-Rabbit secondary antibody (926-32211) (Li-Cor) and visualized using a Li-Cor Odyssey CLx imaging system (Li-Cor).

### Cell viability

A549 cells were treated with drugs resuspended in DMSO at a concentration of 10μM. 24h post treatment, cells were washed in 1 x PBS and cell viability was assessed using CellTiter-Glo Luminescent Cell Viability Assay (Promega). Luminescence read-out was normalized to signal from vehicle treated cells and represented as percentage of viable cells compared to vehicle. Values are an average of triplicate samples and error bars indicate standard deviation.

### RNA-Seq of viral infections and drug treatments

Approximately 1 × 10^6^ A549 cells were infected with SARS-CoV-2. Infections with IAV were performed at a multiplicity of infection of 5 for 9 h in DMEM supplemented with 0.3% BSA, 4.5 g/L D-glucose, 4 mM L-glutamine and 1 μg/ml TPCK-trypsin. Infections with SARS-CoV-2 were performed at an MOI of 0.2 for 24 h in DMEM supplemented with 2% FBS, 4.5 g/L D-glucose, 4 mM L-glutamine, 10 mM Non-Essential Amino Acids, 1 mM Sodium Pyruvate and 10 mM HEPES. Total RNA from infected and mock infected cells was extracted using TRIzol Reagent (Invitrogen) and Direct-zol RNA Miniprep kit (Zymo Research) according to the manufacturer’s instructions and treated with DNase I. RNA-seq libraries of polyadenylated RNA were prepared using the TruSeq RNA Library Prep Kit (Illumina) according to the manufacturer’s instructions. RNA-seq libraries for total ribosomal RNA-depleted RNA were prepared using the TruSeq Stranded Total RNA Library Prep Gold (Illumina) according to the manufacturer’s instructions. cDNA libraries were sequenced using an Illumina NextSeq 500 platform.

### Bioinformatics analyses

Raw reads were aligned to the human genome (hg19) using the RNA-Seq Alignment App on Basespace (Illumina, CA). The top 2000 gene counts with the highest variance were log transformed and Z-score normalized. PCA analysis was performed on the normalized matrix. Differential expression analysis was performed with the Characteristic Direction method on the normalized matrix or with limma on the original count matrix filtered using the method described in (Chen et al., 2016). All differentially expressed genes were submitted for analysis with L1000FWD and Enrichr. Jupyter notebooks from previously published RNA-seq data were generated with BioJupies.

Consensus analysis was performed by counting gene overlap in the top 50 up or down differentially expressed genes based on the consensus L1000FWD and RNA-seq signatures generated from the A549-ACE2 cells. Consensus L1000FWD genes for the drugs were created by enumerating all up and down signatures present in the L1000FWD database associated with those drugs. First the instances of genes appearing in the up or down gene sets were counted, and the cdf of the Z-scored counts was computed to quantify the probability of a given gene being an essential member of signatures for a given drug. Finally, the down probabilities were subtracted from the up probabilities and summed for all drugs resulting in genes ranked by their probable essentiality across all drugs. The top 50 positive (mostly up regulated) and negative (mostly down regulated) genes were taken to represent the consensus genes for these drugs. The RNA-seq signatures from the A549-ACE2 cells were computed by differential expression analysis with limma on the filtered count matrix. The top 50 up and down regulated genes were taken as the signature.

To generate the figure that compares the L1000 differential expression signatures, we obtained all the signatures for the drugs RS-504393, trifluoperazine, chlorpromazine, amlodipine, berbamine, loperamide, and terfenadine from the L1000FWD processed data. Drugs are tested in different cell lines and batches, resulting in 11, 60, 32, 21, 12, 26, and 21 signatures, respectively. We then calculated a representative signature for each drug by averaging the differential expression per gene across the replicates. We then calculated the representative signature for all 4,941 unique drugs contained within the L1000FWD database. We calculated the pairwise similarity between the representative signatures by Pearson correlation. The t-test was applied to compare the pairwise correlation distribution between the 7 candidate drugs and the distribution of all pairwise correlation scores.

### iPSC-derived organoids

Undifferentiated human pluripotent stem cells (hPSCs) were maintained in feeder-free conditions. Briefly, hPSCs were cultured on Matrigel-coated 6-well plates in commercial StemFlex™ Medium (ThermoFisher). The medium was changed every day, and cells were passaged every ~4 days using ReLeSR™ (STEM Technologies). In all normal hPSC cultures, 5 μM Rho-associated protein kinase (ROCK) inhibitor Y-27632 was only added into the culture media when passaging or thawing hPSCs. For pancreatic endocrine organoid differentiation, we adapted previous published protocols (Yang et al., 2020).

### Micro-neutralization assays

293T producer cells were transfected to overexpress SARS-CoV-2, VSV-G, or NiV glycoproteins. Approximately 8 hours post transfection, cells were infected with the VSVΔG-rLuc reporter virus for 2 hours, then washed with DPBS. Two days post infection, supernatants were collected and clarified by centrifugation at 1250 rpm for 5mins. Upon collection, a small batch of VSVΔG-rLuc particles bearing the CoV2pp were then treated with TPCK-treated trypsin (Sigma) at room temperature for 15 minutes prior to inhibition with soybean trypsin inhibitor (SBTI). Particles were aliquoted prior to storage in −80°C to avoid multiple freeze-thaws. To titer these pseudoviruses, 20,000 Vero-CCL81 cells were seeded in a 96 well plate 20-24hrs prior to infection. At 18-22 hours post infection, the infected cells were washed with DPBS, lysed with a passive lysis buffer, and processed for detection of Renilla luciferase. The Cytation3 (BioTek) was used to read luminescence.

---

**Figure S1:**
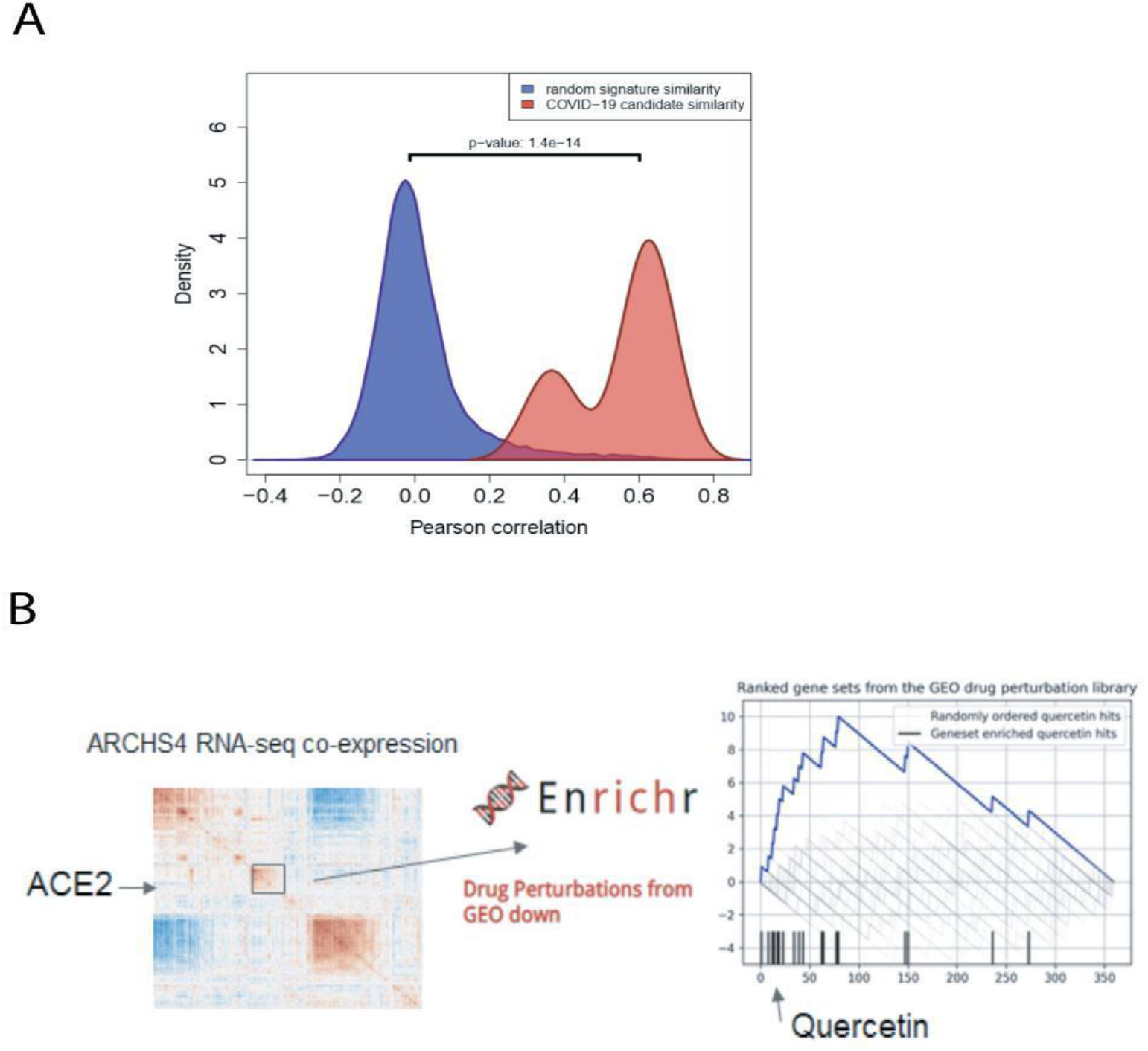
Correlation among predicted drugs in the L1000 space and strategy to predict quercetin. A) Pearson correlation distributions between L1000 L1000FWD signatures for all possible pairs from the 7 predicted (red) and between random pairs of signatures for other L1000 L1000FWD drugs (blue). B) Alternative strategy to predict small molecules for inhibiting SARS-CoV-2. Random walk plot of the ranks of quercetin signatures from the drug perturbation GEO library from CREEDS (right). The ranks are produced by submitting the top 100 most co-expressed genes with the gene ACE2 based on the ARCHS4 co-expression matrix for human RNA-seq data. These 100 genes are submitted to Enrichr to rank drugs against the “Drug Perturbations from GEO down” gene set library.

**Figure S2.**
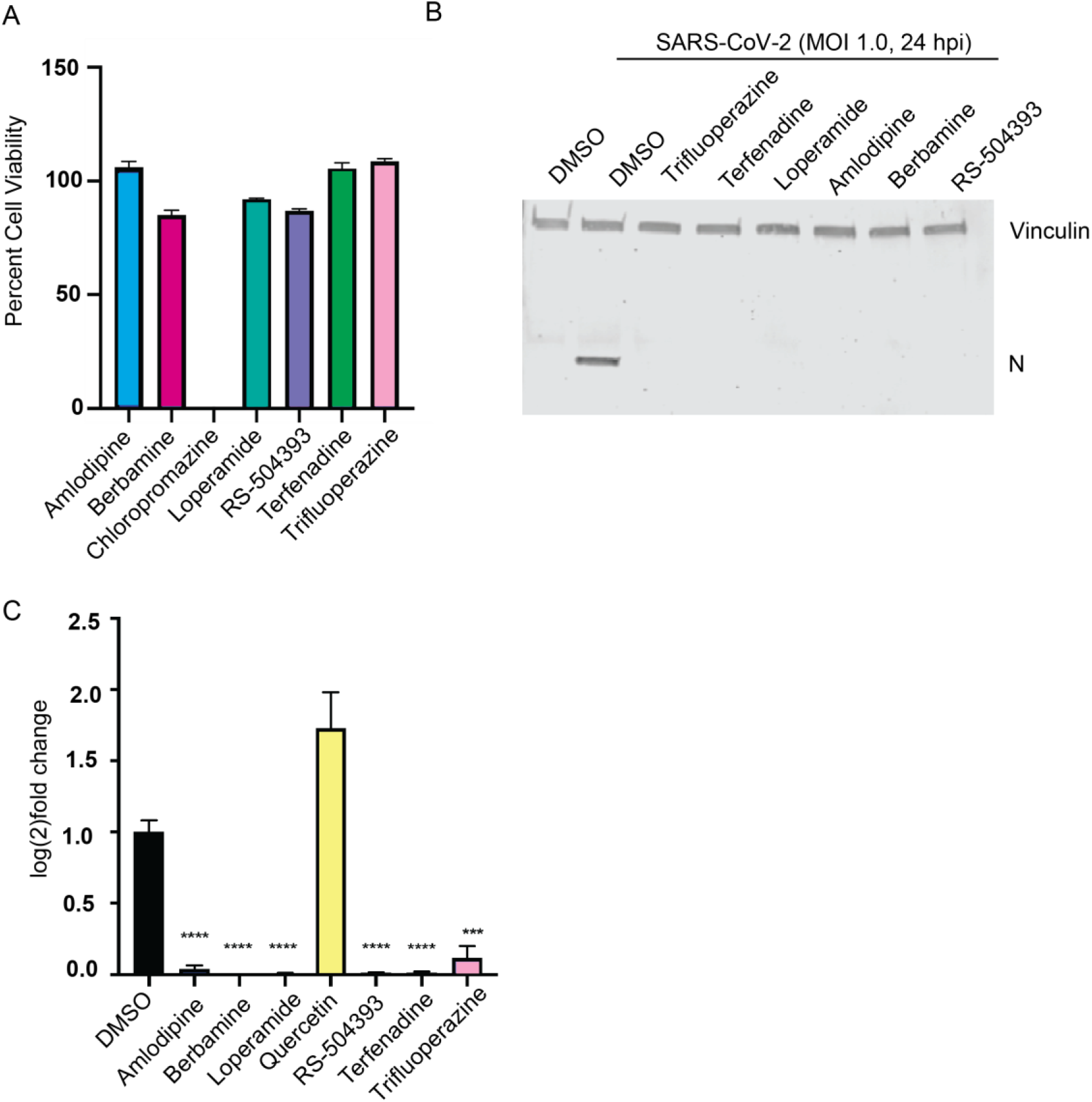
Selected candidate drugs’ cell viability and supporting SARS-CoV-2 inhibition in vitro, Related to Figure 2. A) Percent viability of A549s treated with 10uM of specified drug candidate in DMSO for 24 hours. B) Western blot of ACE2-A549 protein after treatment with specified drug and subsequent SARS-CoV-2 infection at MOI of 0.1 for 24 hours. C) vRNA levels quantified by qRT-PCR analysis of infected Vero-E6 cells after 24 hours of SARS-CoV-2 infection at MOI of 0.01; vRNA levels normalized to human alpha-Tubulin and normalized to DMSO-treated and infected ACE2-A549 RNA. Significance compared to DMSO treated values and determined by unpaired two-tailed student’s t-test, p<0.001 = *** p<0.0001 = ****, n=3 biological replicates for RT-qPCR and cell viability data.

**Figure S3:**
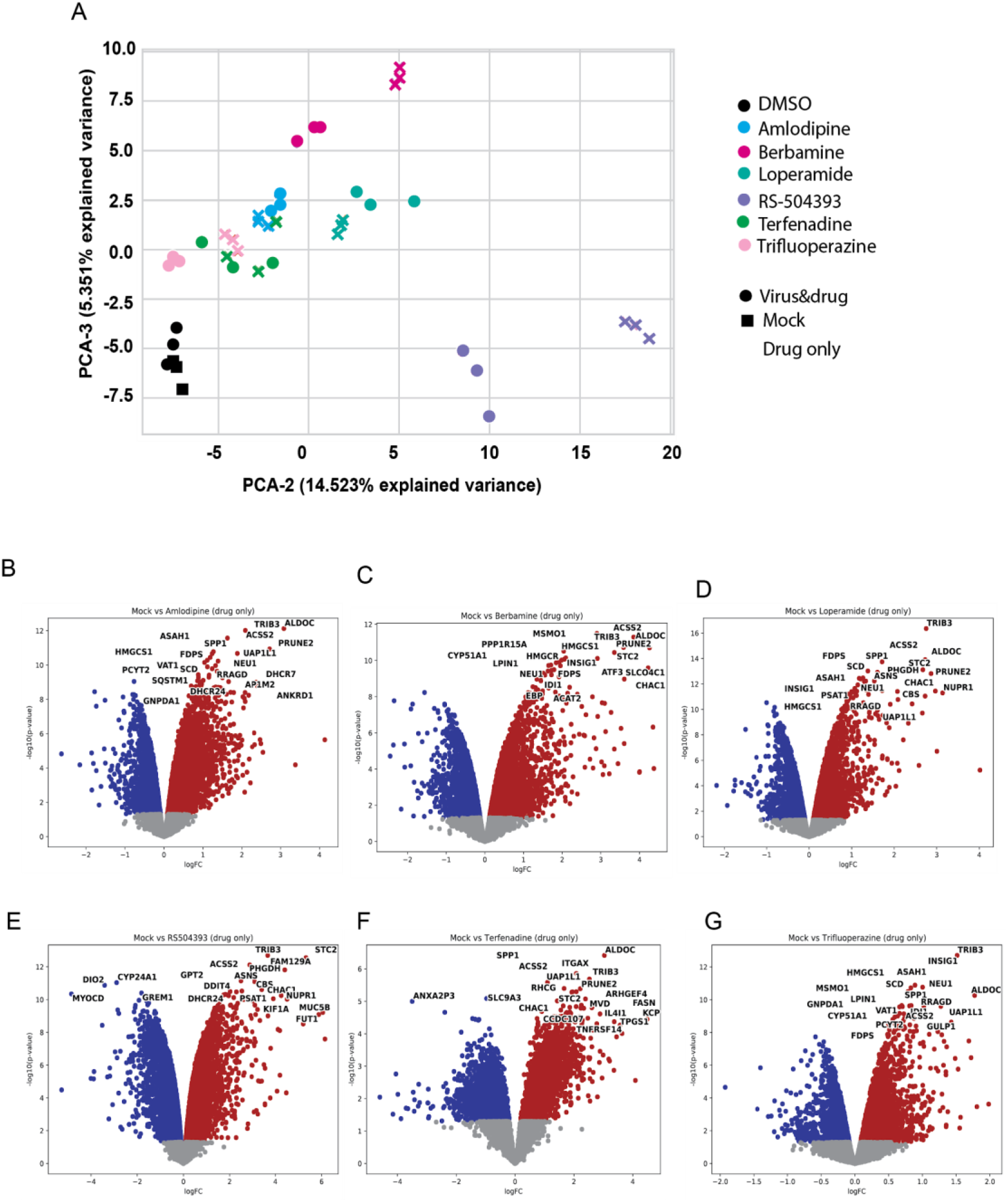
Supporting RNA-seq data analysis from A549^ACE2^ cells, Related to Figure 3. A549^ACE2^ were either mock-infected (MOCK) and treated with DMSO or infected with SARS-CoV-2 in the presence of DMSO or the 6 indicated drugs for 24 hours at MOI 0.1. A) Principal component analysis of the RNA-seq data samples in PC2/PC3 space. D-I) Volcano plots of the A549^ACE2^ populations comparing DMSO- or drug candidate-treated populations without infection, after 24 hours; n=3 biological replicates for RNA-seq data.

**Figure S4:**
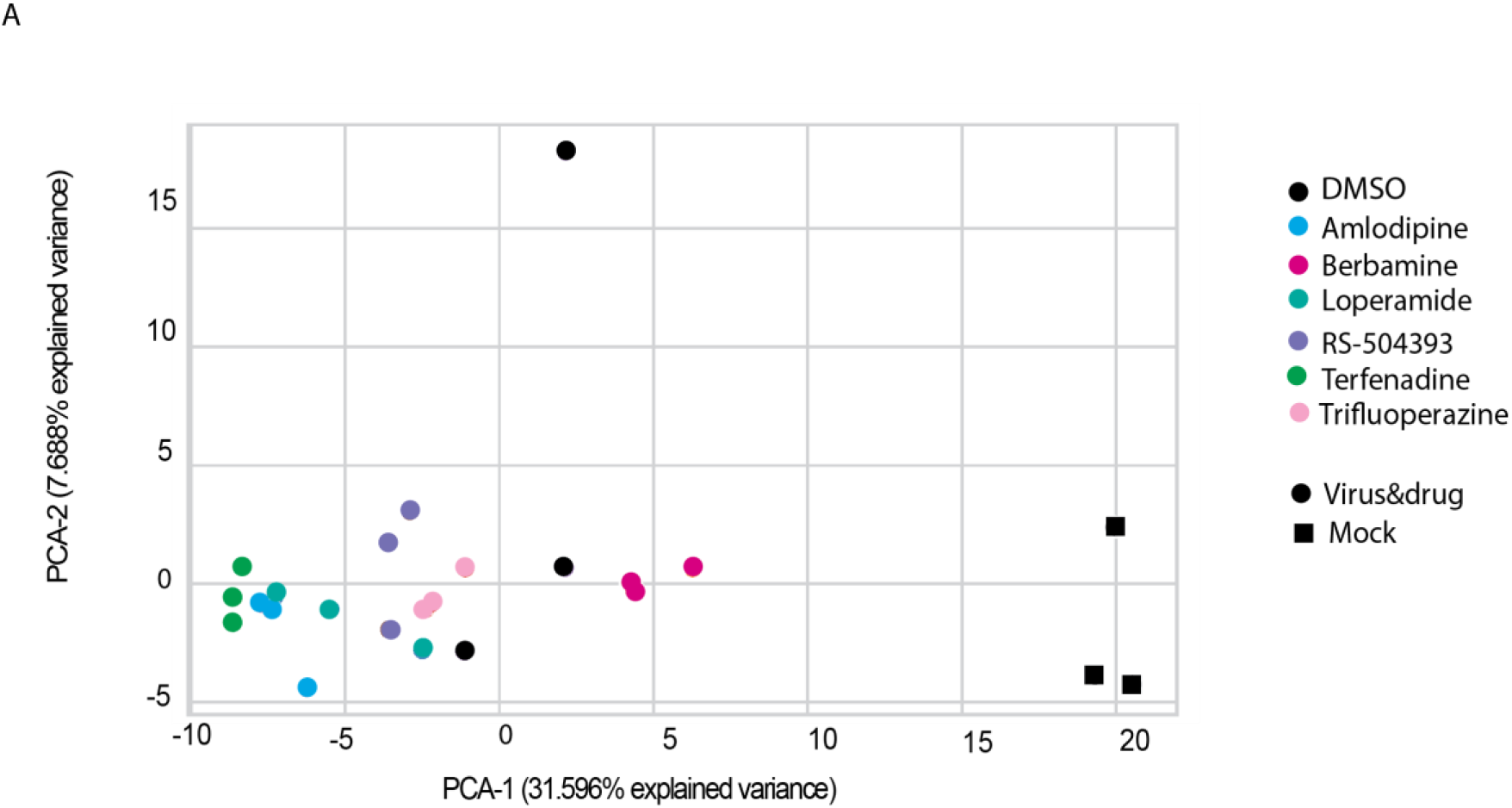
Supporting RNA-seq data analysis from pancreatic endocrine human organoids, Related to Figure 4. Human organoids were either mock-infected (MOCK) and treated with DMSO or infected with SARS-CoV-2 in the presence of DMSO or the 6 indicated drugs for 24 hours at MOI 0.1; n=3 biological replicates. A) PCA Analysis of pancreatic endocrine human organoids in PC2/PC3 space.

## Supporting Online Materials

**Supplementary Tables 1-6** DEGs from A549 cells treated with drug candidates

**Supplementary Tables 7-13** DEGs from A549 cells treated with drug candidates and infected with SARS-CoV-2

**Supplementary Tables 14-20** DEGs from pancreatic organoids treated with drug candidates and infected with SARS-CoV-2

**Supplementary Tables 21-22** BioJupies and Enrichr profiles for each drug condition

